# Comprehensive identification of survival-associated genes for cancers

**DOI:** 10.1101/526285

**Authors:** Hongde Liu, Kun Luo, Huamei Li, Xiao Sun

**Affiliations:** State Key Laboratory of Bioelectronics, School of Biological Science & Medical Engineering, Southeast University, Nanjing 210096, China; Department of Neurosurgery, Xinjiang Evidence-Based Medicine Research Institute, First Affiliated Hospital of Xinjiang Medical University, Urumqi 830054, China

**Keywords:** Prognostic gene, Cancer, Mutation, Diagnosis

## Abstract

Prognostic signature is important in estimating cancer risk, subtyping cancer, and planning treatment. A single gene as a prognostic marker would facilitate the development of a clinical test. Here we showed that the number of prognostic and diagnostic genes differ greatly across cancers. By considering both the survival difference and the fold change of expression in cancer, we revealed the prognostic genes for each cancer and found twenty two genes with both diagnostic and prognostic capacity. The universal prognostic genes (*CDC20, CDCA8, ASPM, ERCC6L*, and *GTSE1*) mainly function in the spindle assembly checkpoint, and show more statistical links to mutated pathways, suggesting that expression of these genes can be altered by mutations from many pathways. Briefly, we systematically identified the prognostic genes and revealed the associations between the prognostic genes and genes mutated in cancer.

**Highlights:**

- The number of possible prognostic and diagnostic genes for cancers;
- A list of independent prognostic genes for each cancer;
- The universal prognostic genes mainly function in the spindle assembly checkpoint;
- Statistical links between mutated pathways and prognostic genes.

## 1 Introduction

Cancers are characterized by distinct patterns of mutation and gene expression, associated with different prognosis.

Prognostic expression signatures have been extensively identified and overall survival exhibits links to diverse biological processes. Uhlen et al summarized that shorter survival is associated with the upregulation of genes related to cell growth and the downregulation of genes related to cellular differentiation.^1^ For breast cancer, the MammaPrint test, including 70 genes, is able to assess the benefit of chemotherapy.^2^ Metastasis is a key factor in short survival. In digestive cancers, besides activation of the mitotic cell cycle, altered expression in the extracellular matrix is linked to poor prognosis.^3–5^ A 64-gene signature is associated with metastasis for non-small cell lung carcinoma (NSCLC).^6^ Inflammatory-related genes are also involved in prognosis in glioblastoma (GBM),^7^ colorectal, and pancreatic cancers.^3,4^ Recently, an estimate of the risk of recurrence of colon cancer was shown to be provided by the total tumor-infiltrating T-cell count and the cytotoxic tumor-infiltrating T-cell count.^8^ In acute myeloid leukemia (AML), the worst overall survival is associated with the expression of multidrug resistant genes.^9^ Also, genes in the tumor microenvironment have a prognostic role for NSCLC.^10^

Cancer-associated mutations have also been extensively studied. Recently, 299 driver mutation genes were revealed.^11^ The study confirmed that microsatellite instability was associated with an improved response to immune checkpoint therapy.^11^ A model estimating the mutation load of 24 genes predicted the response to cancer immunotherapy with anti-CTLA-4 and anti-PD-1 treatments.^12^

The present work focuses on two questions. Firstly, what are the independent prognostic genes for each cancer? The expression signatures in the literature are actually combinations of a set of genes. ^2–7,10,13^ A single gene as a prognostic marker would facilitate the development of a clinical test. Although some independent prognostic genes have been identified, including the genes for *PSA* for prostate cancer,^14^ microRNA-148a for bladder cancer,^15^ telomerase for colorectal cancer,^16^ *KIAA1199* for NSCLC,^17^ and *CTHRC1* for gastric cancer,^18^ a systems-level identification of an independent prognostic gene is still needed. We also noticed that the signature genes differ depending on the criteria. It will be interesting to discover which genes are identified under tighter criteria. Secondly, what are the links between expression of the prognostic genes and the mutations present for a specific cancer? Given that the mutations are the original driving factors for the tumor,^19^ what is the link from the mutations to the altered gene expression, and how does the link determine the prognosis? We will explore the connection.

Here we revealed the independent overall survival-associated genes for each cancer type and identified the genes with capacities of both prognosis and diagnosis. The links between mutation and the prognostic gene expression were also investigated.

## 2 Materials and methods

### 2.1 Datasets

Gene expression data, survival data, and mutation data were retrieved from TCGA project from the initial release of Genomic Data Commons (GDC) in October 2016 using RTCGAToolbox.^20^ A total of 9523 samples across 29 tumor types were downloaded, including 8811 tumor tissues and 712 non-tumor tissues. The abbreviation for cancer type is in supplementary information. Microarray-based gene expression data (gene expression omnibus ID: GSE21501 for pancreatic cancer^21^) were retrieved for validating.

### 2.2 Identification of the prognostic genes

The prognostic genes were identified with a log-rank test in a Kaplan–Meier survival model. In each cancer type, for each gene, patients were classified into two groups, the high-expression group (H) and the low-expression group (L), using the expression median of the gene as a cutoff. In identifying, we considered both survival difference (P[SV]) and the expression change (FC(H/L)) between the two groups. The area under the curve (AUC) of a receiver operating characteristic (ROC) curve and the expression fold change between the cancer (C) and normal (N) tissues (FC(C/N)) were employed to indicate the diagnosis ability.

### 2.3 Regression for the expression of the prognostic genes with the mutation counts

The 40 prognostic genes, which were identified with P[SV]≤10^−6^ and FC[H/L]≥4, and the top 200 frequently mutated genes, were included in this section. Firstly, for each prognostic gene, the dependence between its expression and the mutation counts of the 200 mutated genes were tested with a chi-squared (χ^2^) test for each cancer. The mutated genes with p ≤ 0.001 (χ^2^ test) were included in an enrichment analysis. A cutoff of p ≤ 0.05 was used to find the enriched terms and pathways. Upon satisfaction of those criteria, a link between the prognostic gene and the mutated pathway (terms) was counted. This was done for all cancers to see how many cancer types shared the link. Secondly, we carried out a generalized linear regression of the expression of the prognostic gene with the mutation counts of the top 200 frequently mutated genes for each cancer type. The regression generated a set of parameters indicating the contribution of the mutation in explaining the expression level of the prognostic gene. Only mutated genes with a significant parameter were used to construct a network. More details are in Supplementary Information.

## 3 Results

### 3.1 Prognostic genes differs across cancers

First, we found under the same cutoff of overall survival difference (P[SV]), the number of potential prognostic genes differed between cancer types (Fig. 1A and Fig. S1A). Cancer CHOL, ESCA, STAD, COAD, GBM and PCPG had only a limited number of prognostic genes (Fig. 1A). We assessed the diagnostic genes by calculating the area under the receiver operating characteristic (ROC) curve (AUC) between the cancer and control (normal) data. The cancers also had different numbers of genes at a high AUC value (AUC≥0.9) (Fig. 1A). The result suggests that the number of prognostic and diagnostic genes differs between cancers.

**Figure 1.**
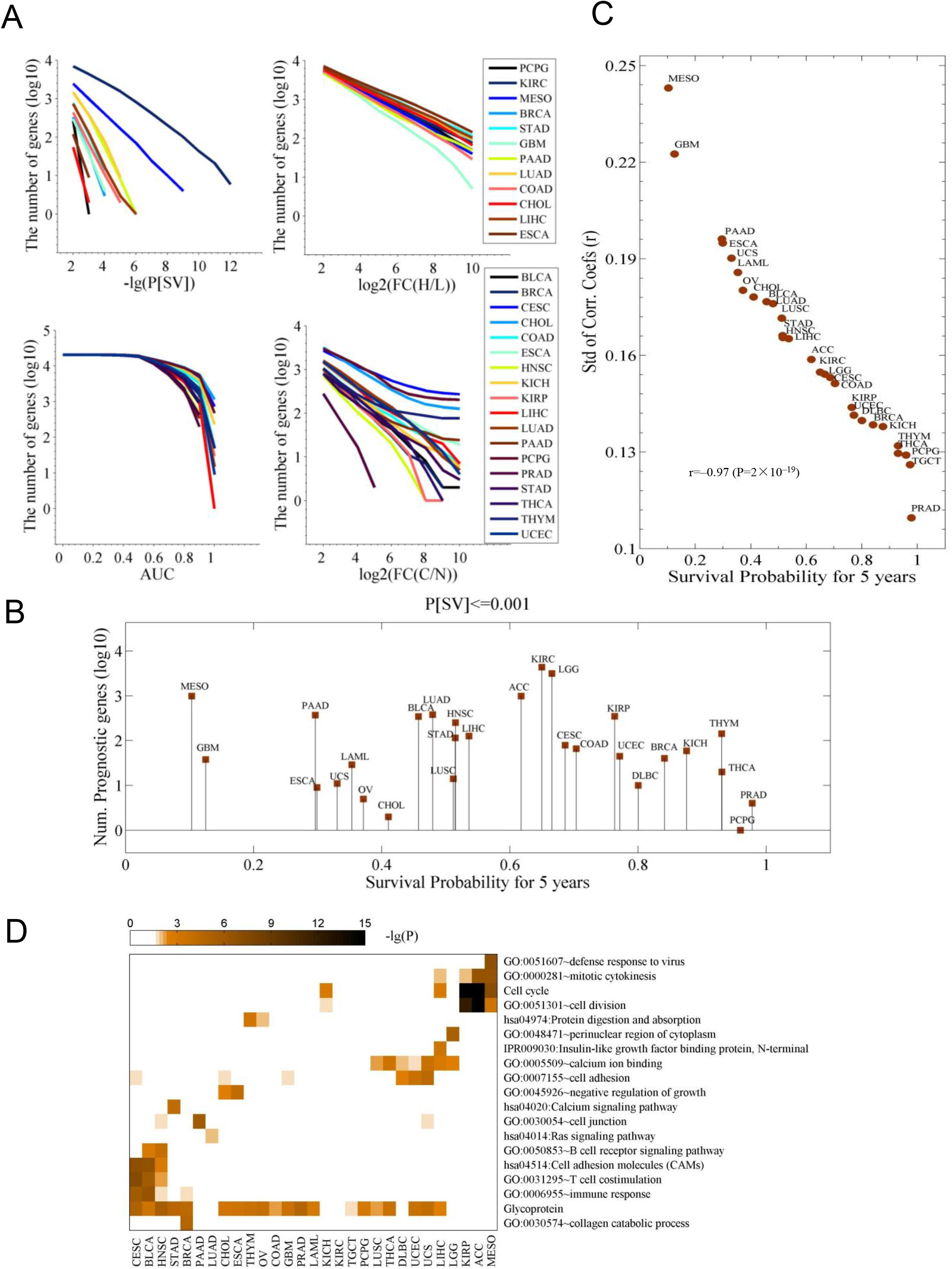
Survival-associated genes differ among cancers. **A**. The number of genes associated with a differential survival (P[SV]), with an expression change in the cancer tissues (FC(H/L)), with a capacity of diagnosis for cancer (AUC), and with an expression change between the control (N) and cancer (C) samples (FC(C/N)), respectively. In each cancer type, for each gene, cancer tissues are divided into high (H) and low (L) expression groups based on the median expression of the gene in the cancer type. P[SV] and FC(H/L) represent survival difference and fold change between the two groups, respectively. FC(C/N) means the fold change between the cancer tissues and normal controls. AUC is the area under receiver operating characteristic (ROC) curves in diagnosing the cancer samples with expression of the gene. **B**. The number of prognostic genes with a significance of P[SV]≤0.001. The data were sorted with the five-year survival probability. **C**. Relationship between survival probability and variation of gene expression among the population. Shown is the five-year survival probability against the standard deviation (Std) of the correlation coefficients of expression profile (20,531 genes) of each pair of patients for each cancer. **D**. The pathways and terms enriched for the genes whose expression was highly correlated to survival time. For each cancer, the top 200 most strongly correlated genes were chosen.

We did not observe a link between the number of prognostic genes and the survival probability (Fig. 1B). We calculated Pearson correlation coefficients (PCCs) of the gene expression of 25,301 genes for each pair of patients for each cancer, and associated the coefficients to the five-year survival probability. Interestingly, the survival probability highly correlated to the standard deviation (Std) of the PCC (r=-0.97) (Fig. 1C).

The pathways and gene ontology (GO) terms that were enriched for the survival-related genes exhibited distinct clusters for the cancer types (Fig. 1D). In KIRP, ACC, and MESO, the genes were enriched for the terms “cell cycle” and “cell division”. CESC, BRCA, STAD, BLCA, and HNSC shared enriched term “T cell costimulation”. LUSC, THCA, DBLC, UCEC, USC, and LIHC had common terms of “cell adhesion”. GBM, LAML, and PAAD were not included in any of the clusters mentioned above. We observed a substantial enrichment in the term “Glycoprotein”. P-glycoprotein was found in the multidrug resistance *(MDR)* phenotype of adult solid tumors.^22^ The enrichment for the genes whose expressions change greatly is shown in Fig. S1B.

### 3.2 Prognostic genes identified

Then, we identified the prognostic genes. We considered two factors to be important. The first was a tight association between the variation of clinical outcome (here, overall survival) and the variation of expression level of the gene, which can be represented by the survival difference (P[SV]) between the high- and low-expression groups. The second was the discriminability of the gene by mRNA level between the two groups. The fold change of gene expression in the high (H) and low (L) groups (FC(H/L)) was used to indicate such discriminability.

First, relaxed criteria of P[SV]≤10^−3^ and FC(H/L)≥2, selecting the top 10 genes for each cancer type, resulted in 236 genes in 29 cancer types (Fig. S2A, Suppl-1). Eight of the 236 genes were in the list identified in literature by P[SV]≤10^−3^ in 17 cancers^1^ (Fig. S2B). The genes include *C1orf88* for ACC, *BCL2L14* for BLCA, *TMEM65* for BRCA, *RBM38* for CESC, *ATP13A3* for CHOL, *ATOH1* for CORD, *ATP1A3* for DLBC and UCS, *GRPEL2* for ESCA, *RARRES2* for GBM, *CHGB* for HNSC, *CLDN3* for KICH, *ATP6V1C2* for KIRH, *HOXD10* for KIRP, *TREML2* for LAML, *ISL2* for LGG, *CDC20* for LIHC, *GTSE1* for LUAD, *PAPPA* for LUSC, *CEP55* for MESO, *DYDC2* for OV, *MYEOV* for PAAD, *KIAA0319* for PRAD, *LBH* for STAD, *CILP* for THCA, *PRKCB* for THYM, and *TP53TG3B* for UCEC (Suppl-1). Very few genes were shared across cancer types, consistent with the literatures.^1,23^ It is notable that the subunits of P- and V-ATPases (such as *ATP13A3, ATP1A3*, and *ATP6V1C2*) were in the list. These genes are responsible for transporting cations across membranes and organelle acidification.^24^

Then, the 236 genes were filtered further with stricter criteria, P[SV]≤10^−6^ and FC(H/L)≥4, resulting in a list of 40 genes (Fig. 2A and Suppl-2). Cancers CHOL, ESCA, TGCT, OV and UCS had no prognostic genes under the criteria (Fig. 2A). The genes *CDC20, CDCA8*, and *CEP55* were prognostic in more than three cancer types. Other genes were specific for particular cancers, such as *MYEOV* for PAAD. Most of the genes had a hazard effect, meaning that high expression of the gene was associated with poorer overall survival (Fig. 2A). The prognostic genes were enriched for the terms of cell cycle, cell division, and cytoskeleton (Fig. S2 C and D). Moreover, in the space of the principal components (PCs) constructed with expression of the 40 genes, the cancer tissues showed clustering patterns and cancer types could be discriminated (Fig. 2B), indicating that the genes represent cancer-type specific survival information.

**Figure 2.**
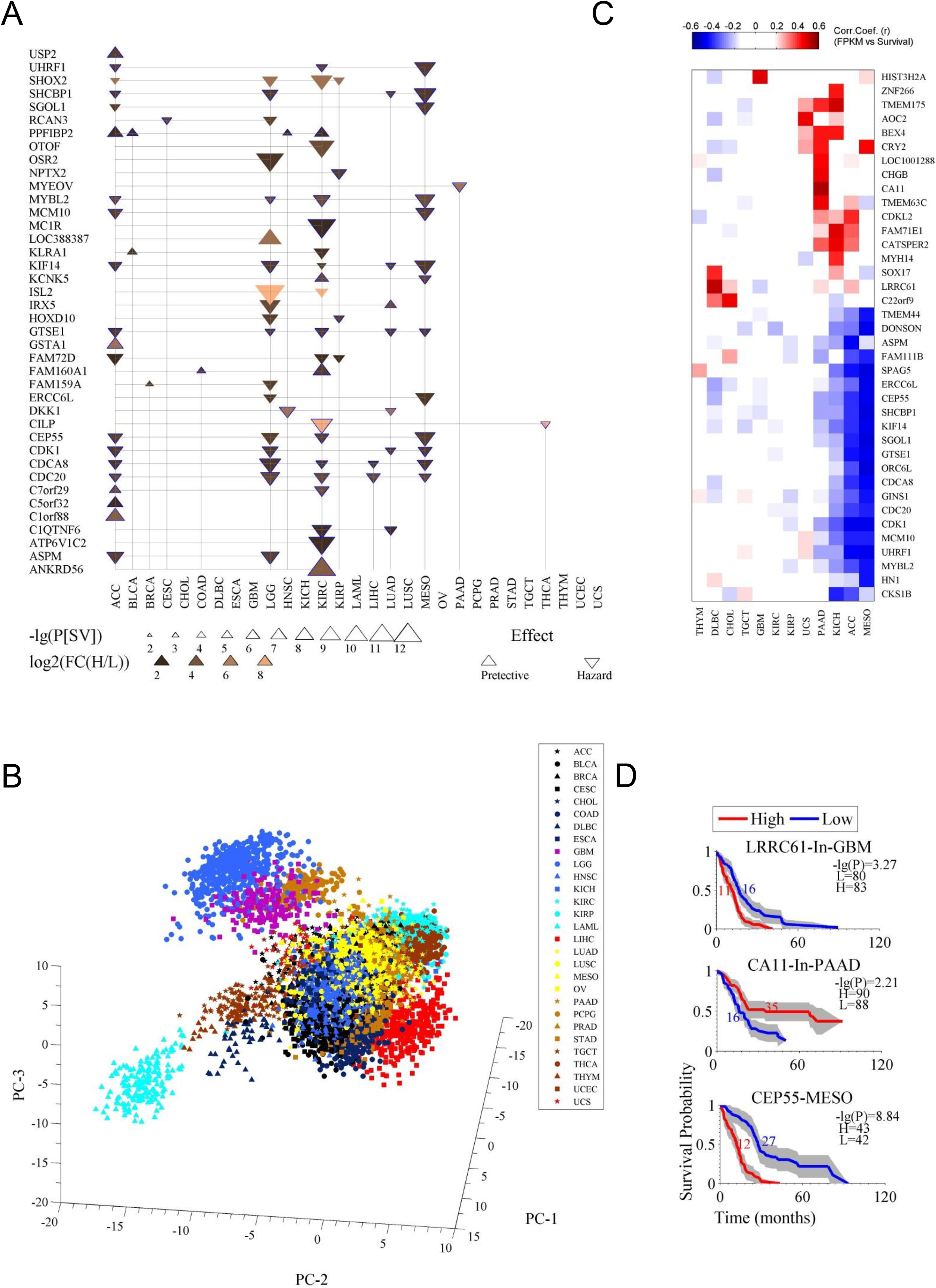
Prognostic genes associated with both survival difference and expression changes in cancer. **A**. The 40 prognostic genes identified. Firstly, with criteria of a survival difference P[SV]≤10^−3^ and fold change of expression FC(H/L)≥2, and by selecting the top ten genes at most for each cancer type, 236 genes were identified in 29 cancer types. Then the 236 genes were further filtered with stricter criteria of P[SV]≤10^−6^ and FC(H/L)≥4. The term “effect” indicates the relationship between gene expression and survival, a downward-pointing triangle means a high expression of the gene corresponds to a poor survival, and an upward-pointing triangle means a high expression to a good survival. **B**. Principal component analysis (PCA) for the gene expression of the 40 prognostic genes in 29 cancer types. **C**. The genes whose expressions highly correlate with survival. The correlation is calculated as a Pearson correlation coefficient (r). **D**. Survival curves for the three prognostic genes.

As few prognostic genes were identified for some cancer types with the strict criteria (Fig. 2A), another strategy, using the Pearson correlation coefficient between survival time and gene expression, was applied to identify optional candidates from the 236 prognostic genes. The result is shown in Fig. 2C. Although some of these genes are not in the list in Fig. 2A, they performed well in indicating survival, for instance, *HIST3H2A* and *LRRC61* for GBM, and *CA11* for PAAD (Fig. 2C and D).

Some of the prognostic genes have been previously identified. For instance, lower expression of *MYBL2* is associated with a favorable overall survival in LGG,^25^ LIHC,^26^ and NSCLC.^27^ High *DKK1* was identified for gastric cancer.^28^ *NPTX2* has been suggested to have prognostic value for GBM,^29^ we found it was moderately significant (p ≤ 0.05) for survival.^30^ We also randomly selected one gene, *MYEOV*, and tested its prognostic performance with microarray-based data for PAAD (Fig. S2E).^21^

### 3.3 Genes with prognostic and diagnostic capacities

We noticed that although some of the genes have prognostic abilities, their average expression levels are even higher in normal tissue than in the low-expression cancer tissues, which leads to confusion in predicting clinical outcome because there is a difficulty in deciding if an unknown tissue is cancer. An example is from gene *DKK1* for LUAD (Fig. S3).

Therefore, we assessed the diagnostic ability of the 236 prognostic genes from two aspects. One was the fold change of gene expression between the cancer (C) and the control (N) tissues (FC(C/N)), and the other was the AUC in diagnosis. With the criteria |log_2_FC(C/N)|≥1.5 and AUC≤0.8, we identified 22 genes (Fig. 3A, Fig. S4, Suppl-3). For *CDC20, CDCA8, CDK1, MYBL2, KIF14, SPAG5*, and *STC2*, their prognostic value has been previously identified. We here confirmed their diagnostic value. For others, we revealed both prognostic and diagnostic values. As shown in Fig. 3B, the expression levels of the genes exhibited a successive increase from the controls, to the low-expression cancer group, and then to the high-expression cancer group, making assessment of both diagnosis and prognosis possible. The genes *CDC20, CDCA8, MYBL2, C1QTNF6, CEP55, CDK1*, and *KIF14* are universal, since they mark multiple types of cancers not only in prognosis but also in diagnosis. *CDC20* can be a prognostic marker for LIHC and KIRC, and can be a diagnostic marker for more than nine types of cancer (Fig. S5). The protein encoded by *CDC20* is required for the full ubiquitin ligase activity of the anaphase promoting complex/cyclosome (APC/C). The spindle assembly checkpoint causes CDC20 to bind to different sites on the APC/C, which alters APC/C substrate specificity.^31^ The product of *CDCA8* is a component of the vertebrate chromosomal passenger complex (CPC), which ensures correct chromosome alignment and segregation and is required for microtubule stabilization and spindle assembly. CDK1 and MYBL2 are central regulators of cell cycle progression. ^33,34^ MYBL2 is a well-known prognostic predictor.^33^ KIF14 is involved in many processes, including chromosome segregation and mitotic spindle formation.^35,36^

**Figure 3.**
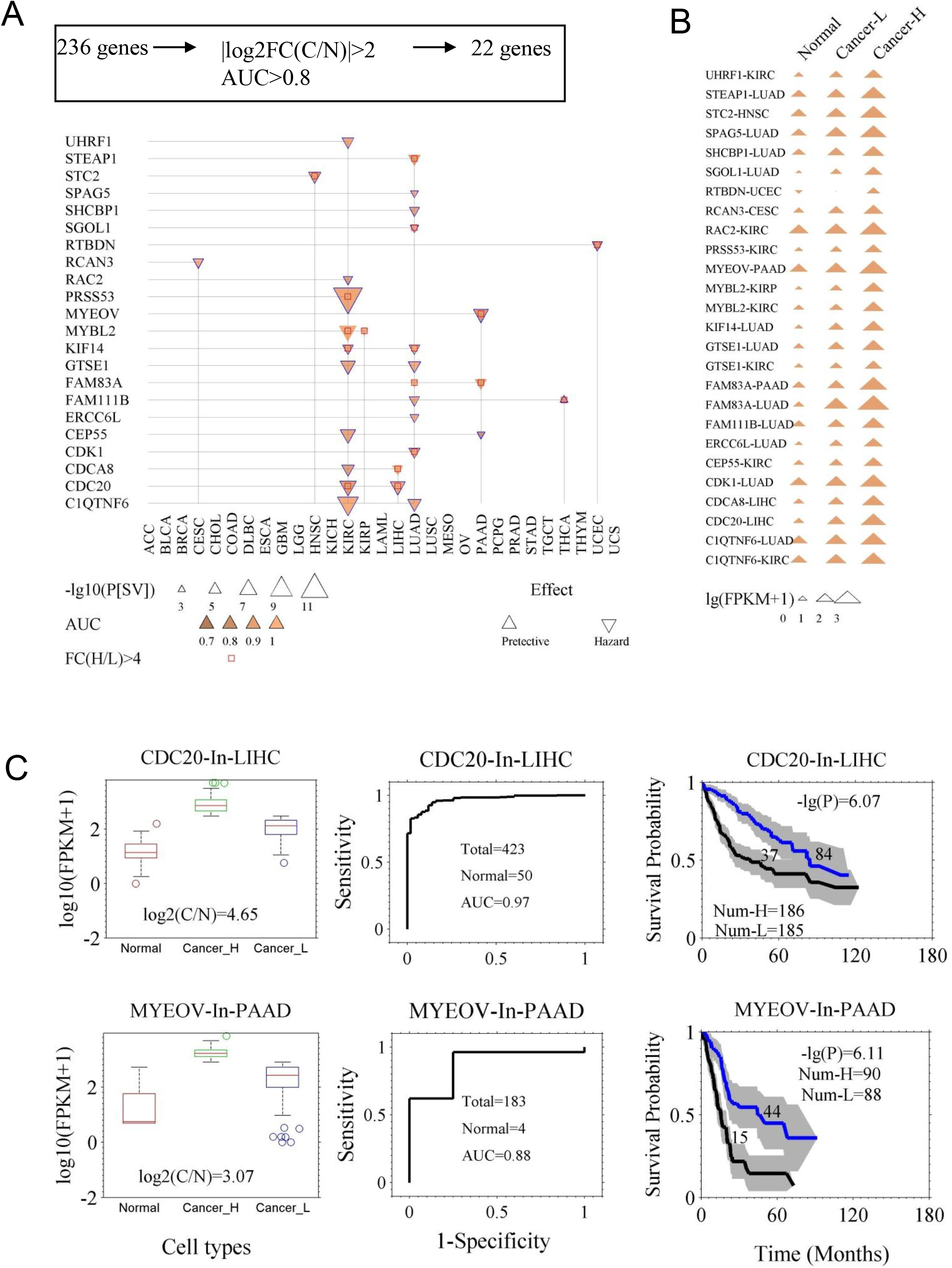
Genes with the capacity for both prognosis and diagnosis. **A**. The 22 genes that are both prognostic and diagnostic. The 236 prognostic genes were further filtered with two new criteria. One was the fold change of expression between the cancer tissues (C) and the normal tissues (N), namely |log_2_(FC[C/N])|≥2. The other was the capacity of differentiating cancer and normal cases, which was assessed by AUC of ROC curve, with the criterion of AUC≥0.8. **B**. The gene expression of the 22 genes in normal tissues, and both low- and high-expression cancer groups (Cancer-L and Cancer-H). **C**. Demonstration of two genes, *CDC20* and *MYEOV*, in prognosis and diagnosis. Left panels, expression levels of the genes in normal and cancer tissues (Cancer-H and Cancer-L). Middle panels, the ROC curves of diagnosis. Right panels, the survival curves of the high- and low-expression groups. The p value of the log-rank test and the number of the groups are indicated.

We also tested the possibility of fifteen immunoregulation-related genes in prognosis as well as diagnosis (Fig. 4). *PVR* was significant for prognosis for KIRC and HNSC. The product of *PVR* is the ligand of TIGIT, which can repress the activity of NK cells.^37^ *CD48* has prognostic value for BRCA (Fig. 4).

**Figure 4.**
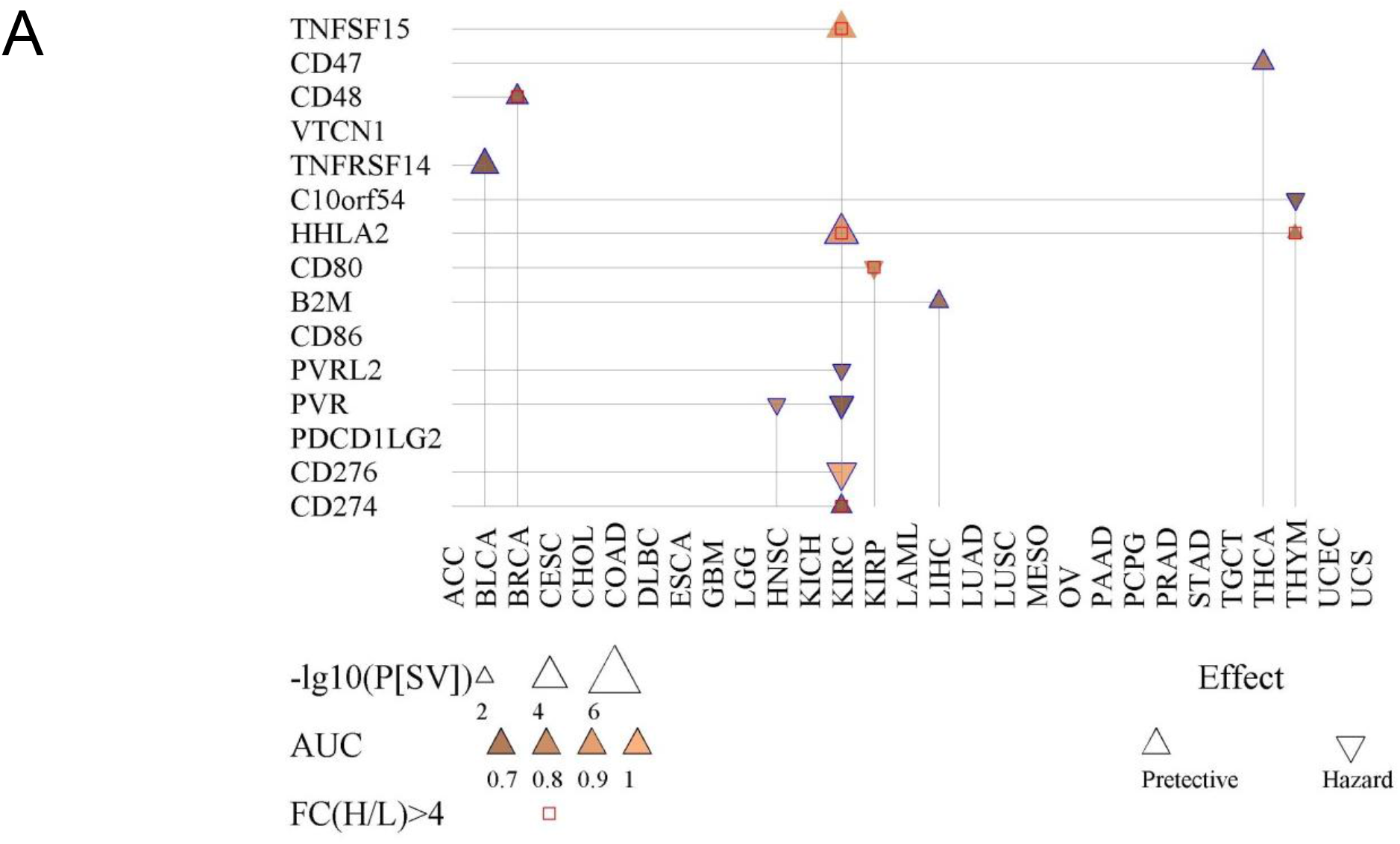
Prognostic and diagnostic value of fifteen immunoregulation-related genes. Shown is the survival difference (P[SV]) between the high- and low-expression groups of each immunoregulation-related gene. The AUC indicates the diagnostic capacity in differentiating cancer tissues.

Briefly, we identified 22 genes that have both prognostic and diagnostic capacities.

### 3.4 Links between mutations and prognostic genes

Finally, we investigated the association between the mutations and the expression of the prognostic genes. The genes with mutations were identified (Fig. S6A and B). The total mutations did not seem to be directly associated with survival (Fig. S6C). We normalized the mutation rate by dividing by gene length, as a longer gene has more chances to mutate.^19^ Interestingly, we found that genes with ~1 mutation per kbp had the lowest expression, in both cancer and control samples (Fig. S6D). Additional analysis is shown in Fig. S7.

We revealed the link between the mutated pathway and the expression of the prognostic genes (Fig. 5A). The prognostic genes could be placed in three classes. In the first class, the gene expression was affected by many mutated pathways in more than five cancer types. These genes were *CDC20, CDCA8*, *ASPM, ERCC6L, KLRA1, KIF14*, *SGOL1*, and *FAM72D*. We noticed *CDC20, ERCC6L, ASPM*, and *CDCA8* are related to anaphase spindle assembly.^31,32,38,39^ The second class included the genes whose expression was affected by only a few pathways, such “Focal adhesion”, the “FoxO and ErbB signaling pathways” and “Carbohydrate digestion and absorption”. These genes included *GTS1, C1orf88, C5orf32, ATP6V1C2, CLIP*, and C1QTNF6. In the last class, links were found to less than three cancer types, showing specificity. The genes *MYEOV, ANKRD56*, and *C7orf29* are connected to mutations in the “Tight junction” and “Long-term potentiation” pathways. Mutations occurring in the PI3K/PI4K domain, methylation-related and central carbon metabolism-related genes showed extensive alteration of the expression levels of the prognostic genes.

**Figure 5.**
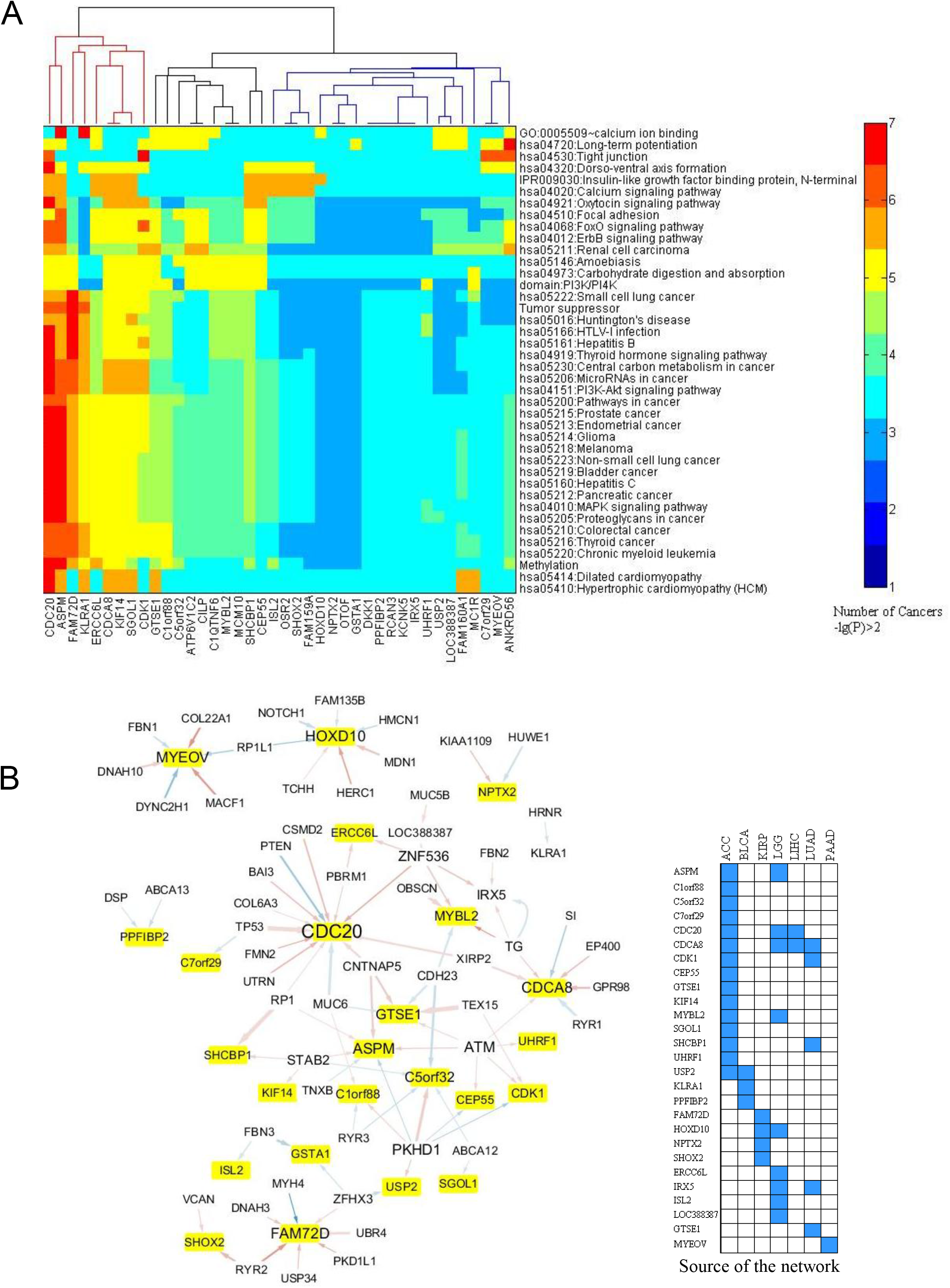
Association between the expression of the prognostic genes and the somatic mutations. **A**. Links between the expression of the prognostic genes and the mutation profile of the top 200 frequently mutated genes. For each prognostic gene, a chi-squared test was employed to test the association between its expression level and the mutation times of the 200 mutated genes in each cancer. The mutated genes with p ≤ 0.001 were included in an enrichment test for GO term, KEGG pathway, InterPro domain, and SMART mode. A cutoff of p ≤ 0.05 was used to find the enriched terms and pathways. If the criteria were satisfied, a link between the prognostic gene and the mutated pathway (term) was counted once. This was done for each cancer type. The heat map here displays the counts for the links within 29 cancer types. **B**. Generalized linear regression of the expression of the prognostic genes with the mutation profile of the top 200 mutated genes. The network displays the regression results after filtering with p < 0.05 in the chi-squared test. In the network, the yellow-marked genes are the prognostic genes and the other genes are the mutated genes. A line from the mutated gene to the prognostic gene indicates that the mutation relates to the expression of the prognostic gene. Blue and red lines mean negative and positive effects, respectively. The line width is proportional to the significance (p value of chi-squared test).

We then tested the relationships between the mutated genes and the prognostic genes (Fig. 5B). *CDC20, CDCA8*, and *ASPM* were associated with a greater number of mutated genes. Mutations in *PKHD1, ATM*, and *ZNF536* were associated with a greater number of prognostic genes.

Frequently mutated genes, including *TP53* and *PTEN*, showed a strong association to *CDC20* expression (Fig. 5B, Fig. 2A). Mutations in *TG* (thyroglobulin), *EP400*, a component of the NuA4 histone acetyltransferase complex, and *SI* (a sucrase-isomaltase enzyme) showed links with *CDCA8*. Mutations in *CNTNAP5*, which encodes a cell adhesion molecule in the nervous system,^40^ and mutations in *ATM*, whose product belongs to the PI3/PI4-kinase family and functions as a cell cycle checkpoint kinase, exhibited a link to the prognostic gene *ASPM. GTSE1* encodes a protein that is involved in p53-induced cell cycle arrest in G2/M phase by interfering with microtubule rearrangements.^41^ We found that mutations in cadherin 23 (*CDH23*), which helps cells stick together, and *TEX15*, which is involved in DNA double-stranded break repair, are linked to *GTSE1* expression. The expression of *MYEOV* is mainly affected by mutations in genes associated with intraflagellar transport (*DYNC2H1*), actin-microtubule interactions and cellular junctions (*MACF1*), myotendinous junctions (*COL22A1*), and calcium-binding microfibrils and glucose homeostasis (*FBN1*).

Briefly, we have provided an insight into the relationships between prognostic genes and genes mutated in cancers.

## 4 Discussion

Prognostic genes are important in estimating low-risk patients, assessing cancer progression, subtyping cancer, and making a proper plan for medical treatment. Here, we reveal the prognostic genes for overall survival for 29 cancers by systematic scanning.

Three points are highlighted. Firstly, the prognostic genes vary greatly among the cancer types. It seems that the more subtypes the cancer has, the fewer prognostic genes. For instance, GBM can be classified into six subgroups (IDH, K27, G34, RTK I and II, and MES). ^42^ Breast tumors have five molecular subtypes (Luminal A, Luminal B, Her2 overexpressing, basal, and normal-like).^43^ ESCA includes four subtypes.^44^ The three cancers had fewer prognostic genes (Fig. 1A-B). In the literature, high levels of intratumor genetic heterogeneity are associated with poorer survival across cancers.^45^ We found a significant association between the intertumor expression heterogeneity and the overall survival (Fig. 1C).

Secondly, the 236 prognostic genes were identified under the criteria of both survival difference and expression change. Expression differentiation between cancer and adjacent normal tissues has been proven to be irrelevant to survival.^46^ We demonstrated it is not appropriate to identify prognostic genes by comparing gene expression between cancer and control samples (*DKK1*, Fig. S3).^46^ We identified 22 genes for both prognosis and diagnosis. High-quality prognostic genes, including *CDC20, ASPM, CDCA8, SGOL1*, and *ERCC6L*, play roles in G2/M processes, such as the spindle assembly checkpoint.^31,32,38,39^ Therefore, the regulation of anaphase of the cell cycle is intimately associated with patient survival.

Thirdly, we associated the mutations and the prognostic genes. Mutations in the PI3K-AKT, ErbB, and FoxO signaling pathways, and in the biological processes of tight junction and methylation, can ubiquitously alter the expression of the prognostic genes. Further relationships were seen between prognostic genes that function in anaphase of the cell cycle, especially *CDC20, CDCA8, ASPM and GTSE1*, and the mutated genes including *TP53, PTEN, ATM, EP400* and *BAI3. PKHD1* mutations link seven prognostic genes (Fig. 5B). Fibrocystin, encoded by *PKHD1* in the liver and kidney, may be involved in cell adhesion, cell repulsion, and the growth and division of cells.^47^

Our results provide a comprehensive prognostic and diagnostic gene list, and reveal the characteristics of prognostic genes of cancer and the statistical association to mutation.

### Contributions

HDL and KL designed the study and wrote the manuscript. HDL and HML did data analysis. KL and XS provided interpretation and discussion. All authors contributed to and approved the final manuscript.

### Declaration of interests

The authors declare that they have no competing interests.

## Acknowledgments

We thank the supports from the National Natural Science Foundation of China (No. 31371339 and No. 81660471) and Key Research & Development Program of Jiangsu Province (BE2016002-3).

## Supplementary Information

### Materials and methods

#### Datasets

Gene expression data, survival data, and mutation data used in this study were retrieved from TCGA project from the initial release of Genomic Data Commons (GDC) in October 2016 (https://portal.gdc.cancer.gov/) using RTCGAToolbox.^1^ A total of 9523 samples across 29 tumor types were downloaded, including 8811 tumor tissues and 712 non-tumor tissues. Microarray-based gene expression data (gene expression omnibus ID: GSE21501 for pancreatic cancer^2^) were retrieved for validating the expression markers for survival.

#### Identification of the prognostic genes

Genes whose expression is associated with a differential overall survival were identified with a log-rank test in a Kaplan–Meier survival model. In each cancer type, for each gene, patients were classified into two groups using the expression median of the gene as a cutoff. The two groups were named the high-expression group (H) and the low-expression group (L), depending on whether the expression level was higher or lower than the median, respectively. The survival difference was tested in the two groups. In identifying, we considered both survival difference and the expression change between the two groups. First, the criteria of a survival difference P[SV]≤10^−3^ and a fold change of expression (FC(H/L))≥2 were applied to 20,531 genes for 29 cancer types, resulting in a list of 236 genes by choosing the top ten genes. Then, stricter criteria, P[SV]≤10^−6^ and FC[H/L]>4 were applied to the 236 genes, resulting in a list of 40 genes. Finally, we were interested in the possibility of the prognostic genes acting as markers to diagnose cancer and normal tissues. Thus, the expression difference between cancer (C) and control (N) tissues was tested. The area under the curve (AUC) of a receiver operating characteristic (ROC) curve and the expression fold change (FC(C/N)) between the cancer and normal tissues were employed to indicate the difference, namely AUC≥0.8 and |log_2_[FC(C/N)]|≥2, resulting in a list of 22 genes.

We also identified prognostic genes another way, by examining the Pearson correlation coefficient between gene expression and survival time among the population. Candidate genes were those with a large positive or negative correlation.

#### Analysis of association between immunoregulation-related gene expression and survival time

Fifteen immunoregulation-related genes were considered, including *PDL-1, PDL-2*, and *PVR*. PDL-1 and PDL-2 are ligands of programmed death-1 (PD-1), which is involved in the inhibition of T cells,^3^ and PVR is a ligand of TIGIT. The binding of the two inhibits NK cell cytotoxicity.^4^

#### Analysis of the mutations in cancers

For each gene, mutations were counted in all cancers. A mutation rate (frequency) of the gene is the ratio of the total mutations in the gene to the number of all patients in all cancers. The top 200 frequently mutated genes were used in a regression model for the expression of the prognostic genes to examine what kind of mutations contributed to the expression levels of the prognostic genes (see below). Given that the mutation count is related to both gene length and gene expression,^5,6^ the mutation rate was normalized by dividing the rate by gene length [Kbp].

We investigated the relationship between the mutation counts and the survival time. The survival time was truncated at a survival probability of 60%. We also tested the relationship between the normalized mutation rate and the expression level.

#### Regression for the expression of the prognostic genes with the mutation counts

The 40 prognostic genes, which were identified with P[SV]≤10^−6^ and FC[H/L]≥4, and the top 200 frequently mutated genes, were included in this section of analysis. Firstly, the prognostic genes were aligned to the pathways enriched for the mutated genes and GO terms. For each prognostic gene, the dependence between its expression and the mutation counts of the 200 mutated genes were tested with a chi-squared (χ^2^) test for each cancer. The mutated genes with p ≤ 0.001 (χ^2^ test) were included in an enrichment analysis on GO terms, KEGG pathways, InterPro domains, and SMART modes, based on a hypergeometric distribution or Fisher’s exact test. A cutoff of p ≤ 0.05 was used to find the enriched terms and pathways. Upon satisfaction of those criteria, a link between the prognostic gene and the mutated pathway (terms) was counted. This was done for all cancers to see how many cancer types shared the link.

Secondly, we carried out a generalized linear regression of the expression of the prognostic gene with the mutation counts of the top 200 frequently mutated genes for each cancer type. The regression generated a set of parameters indicating the contribution of the mutation in explaining the expression level of the prognostic gene. Only mutated genes with a significant parameter were used to construct a network.

#### Enrichment analysis

The enrichment analysis for the 236 prognostic genes and the top 200 frequently mutated genes was carried out with DAVID.^7^

#### Other analysis in the work

The activation status of pathways was assessed with SPIA.^8^ Survival analysis and the survival comparison were carried out by an inhouse Matlab program (Suppl-4).

## Abbreviation for cancer type

LAML,: Acute Myeloid Leukemia;
ACC,: Adrenocortical carcinoma;
BLCA,: Bladder Urothelial Carcinoma;
LGG,: Brain Lower Grade Glioma;
BRCA,: Breast invasive carcinoma;
CESC,: Cervical squamous cell carcinoma and endocervical adenocarcinoma;
CHOL,: Cholangiocarcinoma;
LCML,: Chronic Myelogenous Leukemia;
COAD,: Colon adenocarcinoma;
ESCA,: Esophageal carcinoma;
GBM,: Glioblastoma multiforme;
HNSC,: Head and Neck squamous cell carcinoma;
KICH,: Kidney Chromophobe;
KIRC,: Kidney renal clear cell carcinoma;
KIRP,: Kidney renal papillary cell carcinoma;
LIHC,: Liver hepatocellular carcinoma;
LUAD,: Lung adenocarcinoma;
LUSC,: Lung squamous cell carcinoma;
DLBC,: Lymphoid Neoplasm Diffuse Large B-cell Lymphoma;
MESO,: Mesothelioma;
OV,: Ovarian serous cystadenocarcinoma;
PAAD,: Pancreatic adenocarcinoma;
PCPG,: Pheochromocytoma and Paraganglioma;
PRAD,: Prostate adenocarcinoma;
STAD,: Stomach adenocarcinoma;
TGCT,: Testicular Germ Cell Tumors;
THYM,: Thymoma;
THCA,: Thyroid carcinoma;
UCS,: Uterine Carcinosarcoma;
UCEC,: Uterine Corpus Endometrial Carcinoma.

---

**Figure S1.**
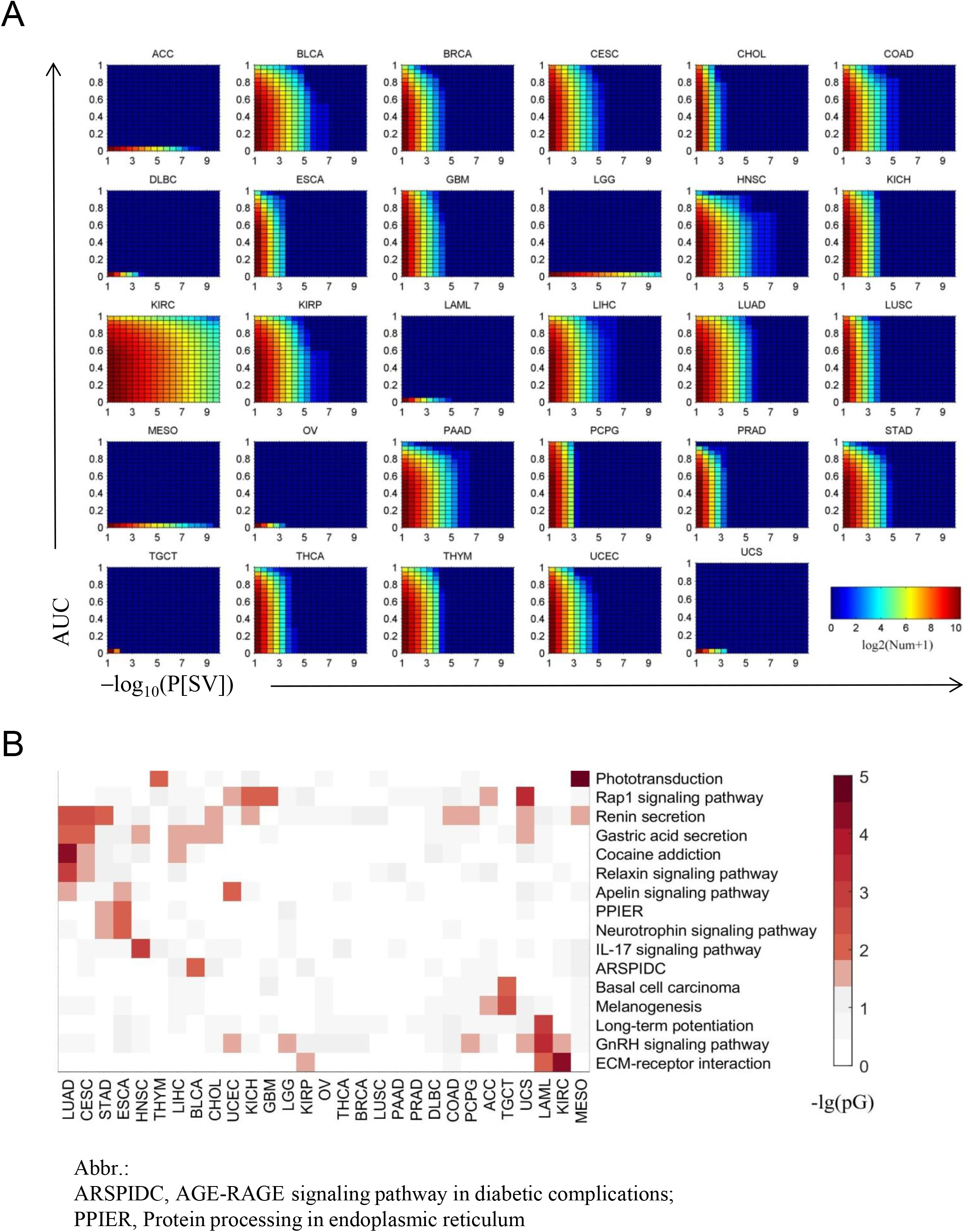
Cancers exhibit a great difference in the number of the genes that are associated with survival difference P[SV] and have a diagnostic value (AUC). **A**. Gene numbers at different cutoffs of both survival difference and differential expression between cancer and control. For each gene in each cancer, P[SV] was calculated with a log-rank test in the high- and low-expression groups that were obtained by dividing the cancer tissues by the median expression of the gene. AUC is the area under the ROC curves in differentiating the cancer and the control with the gene expression. **B**. The pathways that are significantly enriched for the genes whose expression changes greatly among cancer tissues (FC(H/L)≥2). The analysis was done with the SPIA tool.

**Figure S2.**
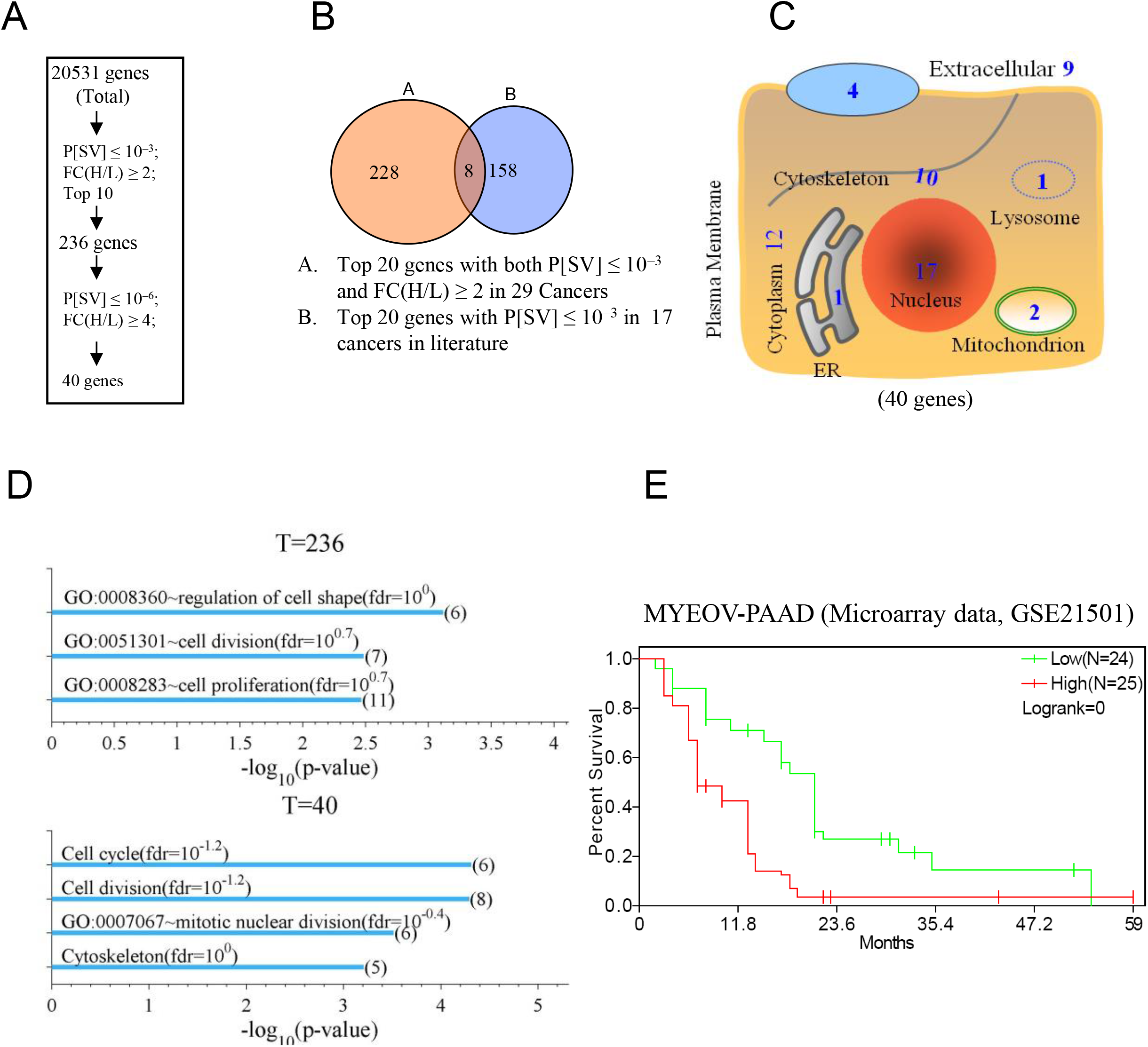
Forty prognostic genes identified by the criteria P[SV]≤10^−6^ and FC[H/L]≥4. **A**. The identification flow chart. P[SV] is from the log-rank test in the high- and low-expression groups that were obtained by dividing the cancer tissues by the gene’s median expression. **B**. Venn diagram showing the number of the overlapping genes in our study (A) and in the literature (B) (Uhlen M. *et al*. 2017, Science 357). The overlapping genes are *DKK1, EMP1, EPS8, FAM83A, MET, MGAT4A, MYEOV*, and *PGK1*. **C**. Subcellular location of the forty prognostic genes. **D**. An enrichment analysis for the identified 236 and 40 prognostic genes with the DAVID tool for KEGG pathways and GO terms. **E**. The survival curves of the *MYEOV* high- and low-expression cancer groups in PAAD using microarray data (GEO ID, GSE21501).

**Figure S3.**
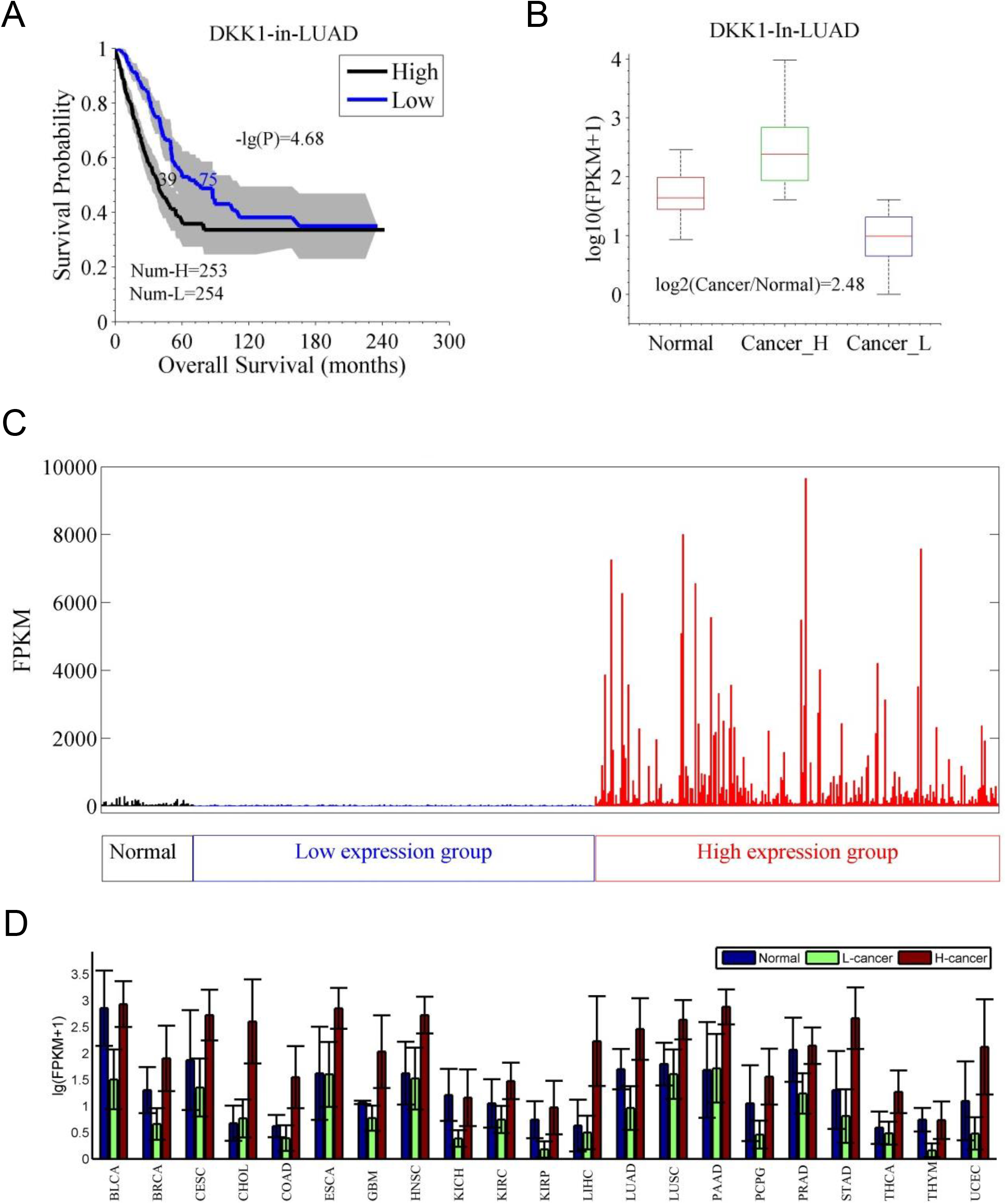
Sample to demonstrate that a prognostic gene is not necessarily suitable for discriminating cancer tissues from normal tissues. **A**. The survival curves of the *DKK1* high- and low-expression cancer groups. **B**. Boxplot of the expression of *DKK1* in LUAD tissues and normal tissues. LUAD tissues are divided into high- and low-expression groups based on the median expression. **C**. Expression of *DKK1* in each LUAD tissue. **D**. Average expression of *DKK1* in 21 cancers and controls.

**Figure S4.**
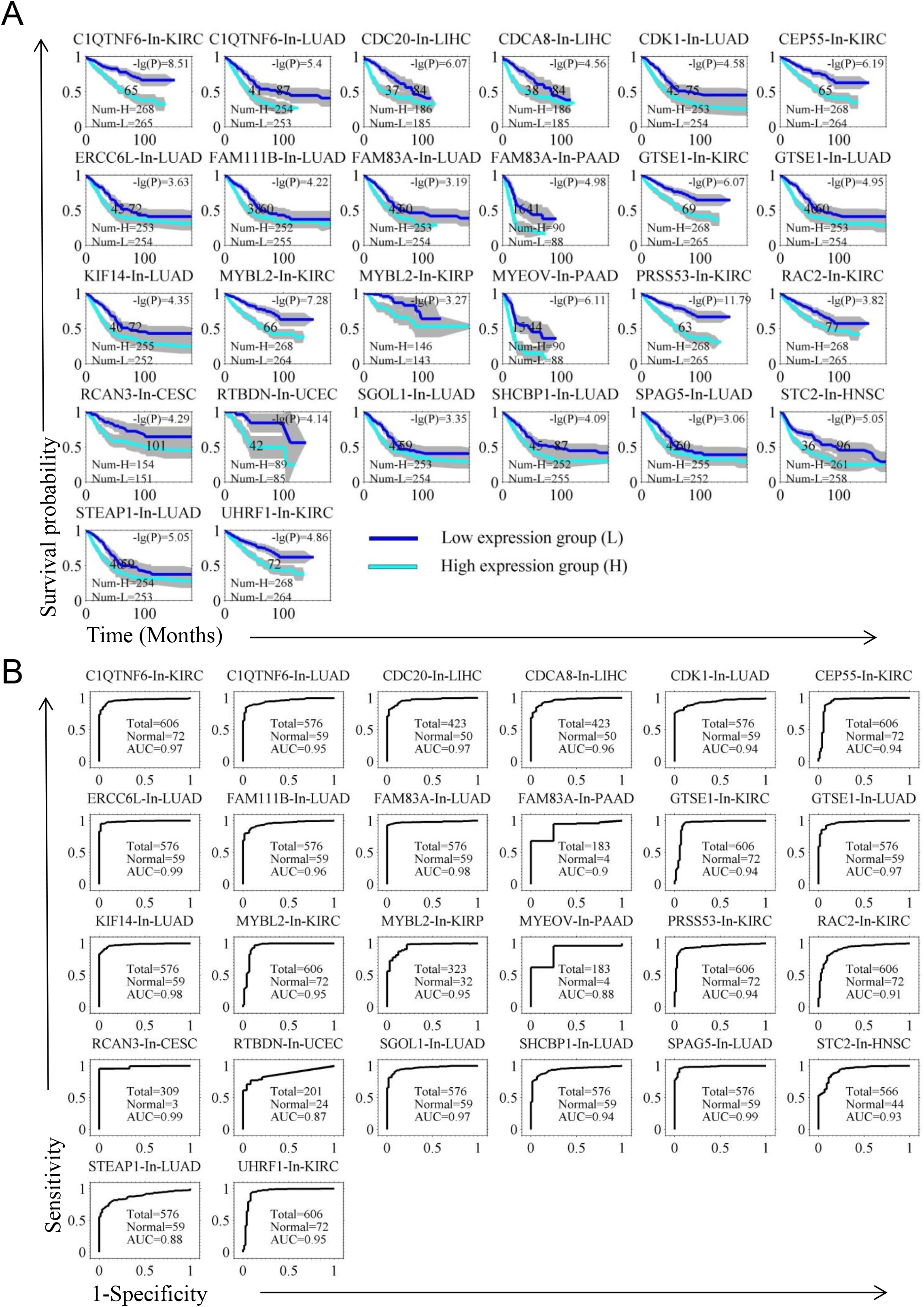
Survival curves and ROC curves to show the capacity of the genes in both prognosis and diagnosis in specific cancers. **A**. The survival curves of the high- and low-expression groups. The difference significance P[SV] and the median survival time are indicated. **B**. ROC curves of diagnosis.

**Figure S5.**
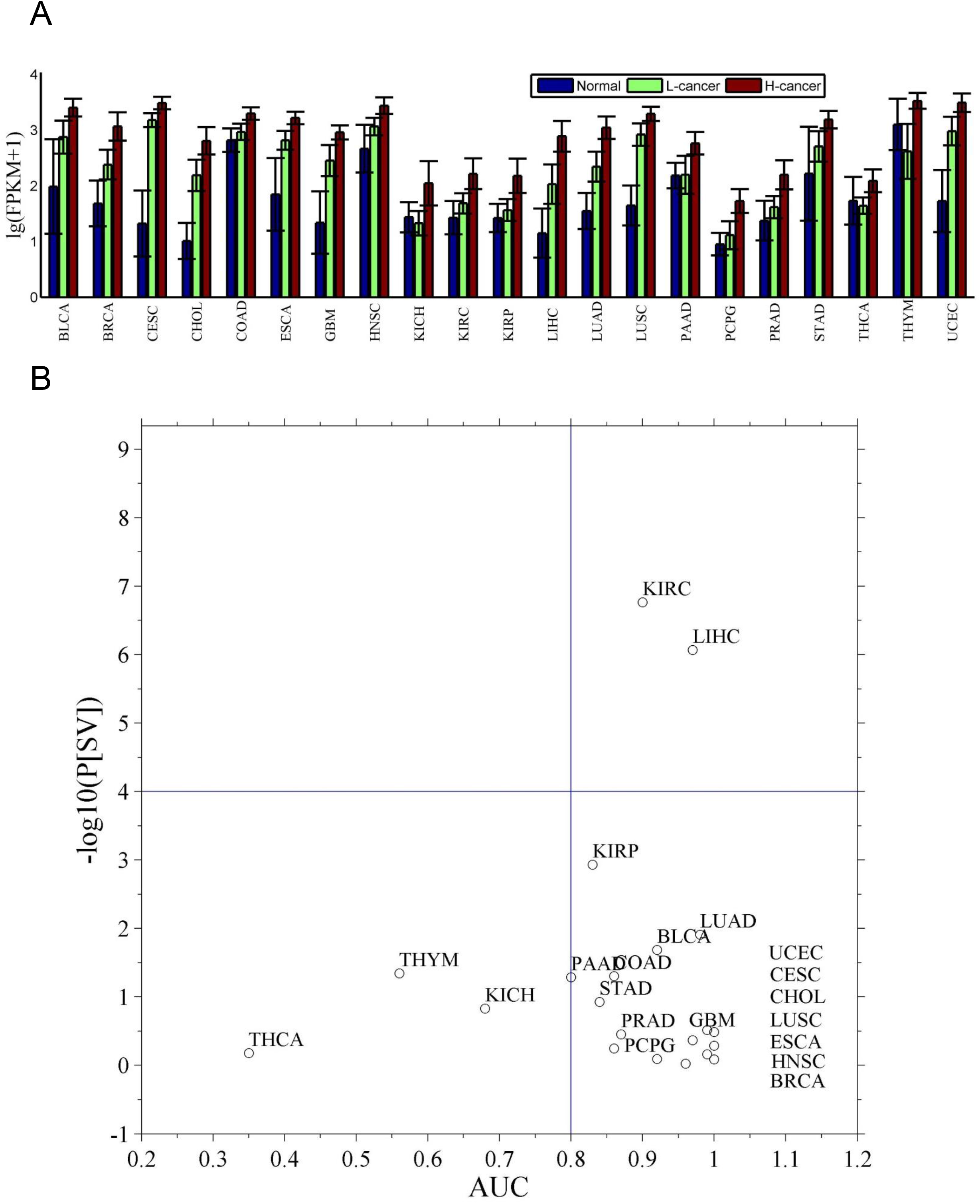
Characteristics of *CDC20* in prognosis and diagnosis in 21 cancers. **A**. The boxplots indicate gene expression of *CDC20* in normal tissues, and the high- and low-cancer groups. **B**. Plot of the prognostic and diagnostic values of *CDC20* in cancers.

**Figure S6.**
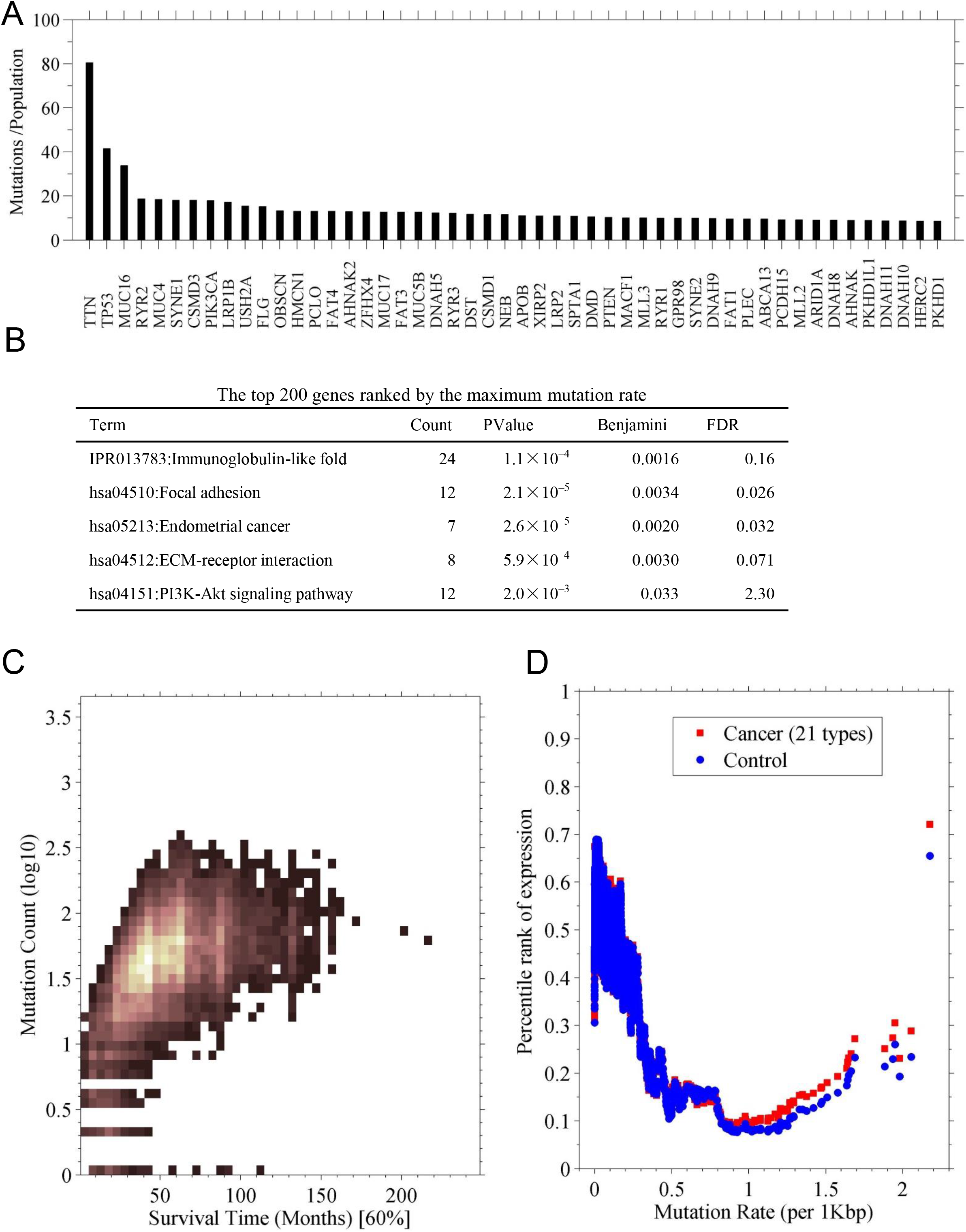
Most frequently mutated genes in cancers. **A**. Shown are the top 40 mutated genes for all cancer types. The vertical axis represents the mutation rate, namely a rate of total number of mutations occurring in the gene to the number of all patients in all cancer types. **B**. An enrichment analysis for the top 200 frequently mutated genes. **C**. Survival time of the patients with the mutations in each gene. The survival time is truncated at a survival probability of 60%; each patient that has a mutation in the gene was counted. Shown is a heat map indicating the distribution of the survival time against the mutation counts. Lightness is proportional to the number of genes. **D**. Relationship between the expression level and the mutation rate for each gene. The mutation rate was normalized by dividing the rate by gene length [Kbp]. The expression data (FPKM) were normalized into percentile in each sample (patient). For each gene, its expression percentile was plotted to its mutation rate per base pair.

**Figure S7.**
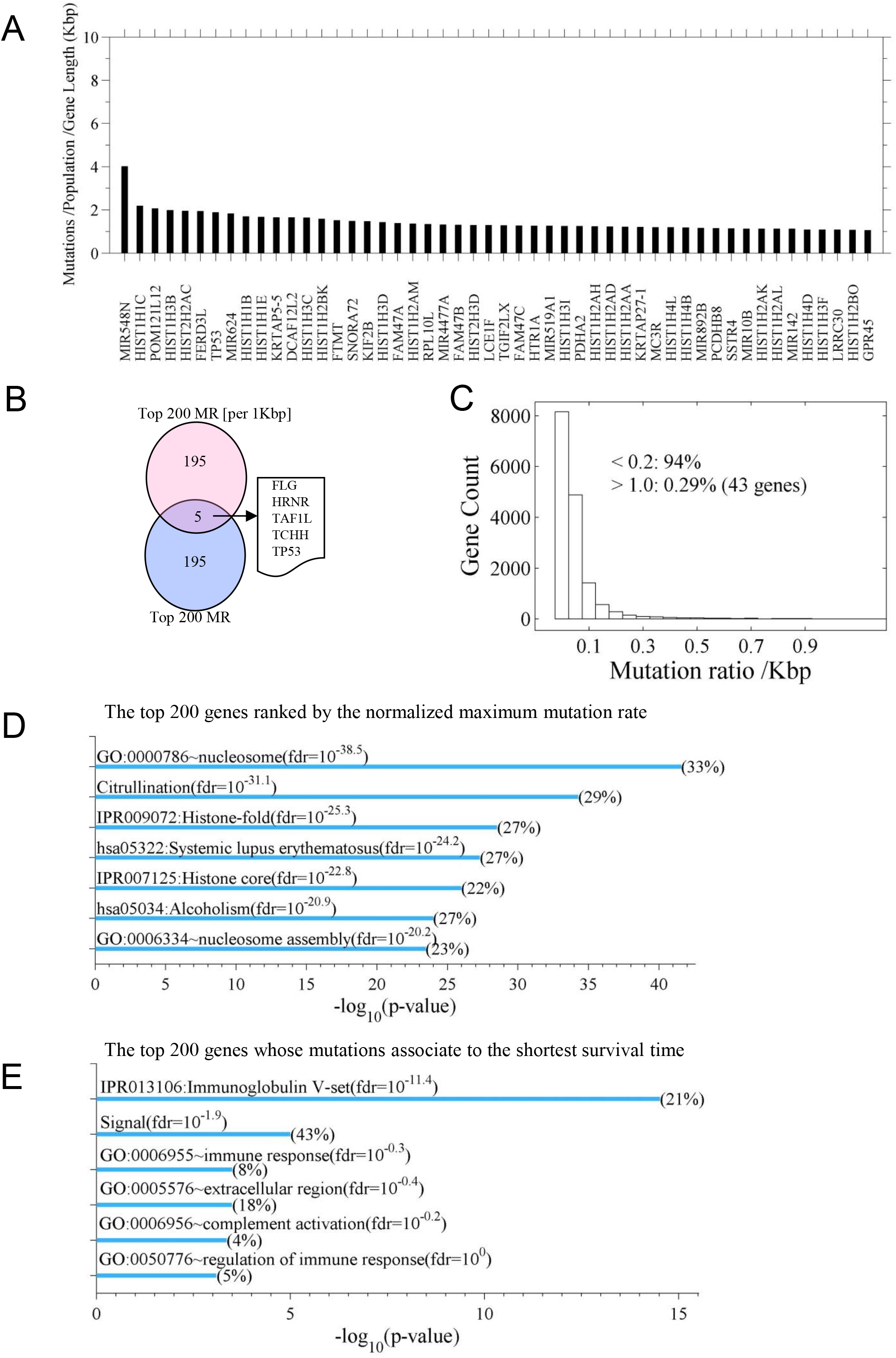
**A**. Top 40 mutated genes for all cancer types. The mutation rate was normalized by dividing by gene length [Kbp]. **B**. Venn diagram showing the overlapping genes between the top 200 genes ranked by the maximum mutation rate before and after the normalization by gene length. **C**. Distribution of the normalized mutation rate. **D**. An enrichment analysis for the top 200 genes ranked by the normalized maximum mutation rate. **E**. An enrichment analysis for the top 200 genes whose mutations associate to the shortest survivals (from Fig. S6C).

## Supplementary Files

### Suppl-1

The prognostic genes identified at criteria of P[SV]≤10^−3^ and FC(H/L)≥2, selecting the top ten genes for each cancer type. There were a total of 236 genes in 29 cancer types. Here, in each cancer type, for each gene, patients are classified into high (H) and low (L) expression groups using the expression median of the gene as a cutoff. The survival difference (P[SV]) was tested between the two groups with a log-rank test. FC(H/L) means the fold change of average expression between the two groups. In the files, chi-square is the value of χ^2^ in the log-rank test, T-middle-H and T-middle-L are the time at 50% survival probability for the high- and low-expression groups, respectively, and mean-H and mean-L are the mean expression in the high- and low-expression groups, respectively. The gene symbol and cell type are the official symbol of the prognostic gene and the corresponding cancer type, respectively.

### Suppl-2

The prognostic genes identified at criteria of P[SV]≤10^−6^ and FC(H/L)≥4. In the file, the term “Effective” indicates the relationship between gene expression and survival, a downward-pointing triangle means a high expression of the gene corresponds to poor survival, and an upward-pointing triangle means a high expression of the gene corresponds to good survival. The other settings are the same as in the Suppl-1 file.

### Suppl-3

The genes that have both prognostic and diagnostic values, with criterion of P[SV] ≤ 10^−4^, indicating the survival difference between the high and low expression groups, and AUC ≥ 4, indicating the capacity of the diagnosing cancer. The term log_2_(FC(C/N)) is the fold change of average expression between the cancer (C) and the control (N). Other settings are same as in Suppl-1 file.

### Suppl-4

The programs (in Matlab) that used in the analysis;

“Survival_Analysis_OKAY.m”, for survival analysis based on Kaplan-Meier survival model. “SR_chi_test_bigdata.m”, for survival difference test, namely log-rank test. “plot_Survival_Anlysis”, for plotting the survival curves.

## References

1 Uhlen, M. et al. A pathology atlas of the human cancer transcriptome. Science 357, doi:10.1126/science.aan2507 (2017).

2 Cardoso, F. et al. 70-Gene Signature as an Aid to Treatment Decisions in Early-Stage Breast Cancer. The New England journal of medicine 375, 717–729, doi:10.1056/NEJMoa1602253 (2016).

3 Birnbaum, D. J. et al. A 25-gene classifier predicts overall survival in resectable pancreatic cancer. BMC medicine 15, 170, doi:10.1186/s12916-017-0936-z (2017).

4 Barrier, A. et al. Colon cancer prognosis prediction by gene expression profiling. Oncogene 24, 6155–6164, doi:10.1038/sj.onc.1208984 (2005).

5 Nault, J. C. et al. A hepatocellular carcinoma 5-gene score associated with survival of patients after liver resection. Gastroenterology 145, 176–187, doi:10.1053/j.gastro.2013.03.051 (2013).

6 Lu, Y. et al. A gene expression signature predicts survival of patients with stage I non-small cell lung cancer. PLoS medicine 3, e467, doi:10.1371/journal.pmed.0030467 (2006).

7 Kim, Y. W. et al. Identification of prognostic gene signatures of glioblastoma: a study based on TCGA data analysis. Neuro-oncology 15, 829–839, doi:10.1093/neuonc/not024 (2013).

8 Pages, F. et al. International validation of the consensus Immunoscore for the classification of colon cancer: a prognostic and accuracy study. Lancet 391, 2128–2139, doi:10.1016/S0140-6736(18)30789-X (2018).

9 Wilson, C. S. et al. Gene expression profiling of adult acute myeloid leukemia identifies novel biologic clusters for risk classification and outcome prediction. Blood 108, 685–696, doi:10.1182/blood-2004-12-4633 (2006).

10 Navab, R. et al. Prognostic gene-expression signature of carcinoma-associated fibroblasts in non-small cell lung cancer. Proceedings of the National Academy of Sciences of the United States of America 108, 7160–7165, doi:10.1073/pnas.1014506108 (2011).

11 Bailey, M. H. et al. Comprehensive Characterization of Cancer Driver Genes and Mutations. Cell 174, 1034–1035, doi:10.1016/j.cell.2018.07.034 (2018).

12 Lyu, G. Y., Yeh, Y. H., Yeh, Y. C. & Wang, Y. C. Mutation load estimation model as a predictor of the response to cancer immunotherapy. NPJ genomic medicine 3, 12, doi:10.1038/s41525-018-0051-x (2018).

13 Abdul Aziz, N. A. et al. A 19-Gene expression signature as a predictor of survival in colorectal cancer. BMC medical genomics 9, 58, doi:10.1186/s12920-016-0218-1 (2016).

14 Palmberg, C., Koivisto, P., Visakorpi, T. & Tammela, T. L. PSA decline is an independent prognostic marker in hormonally treated prostate cancer. European urology 36, 191–196, doi:10.1159/000067996 (1999).

15 Ma, L., Xu, Z., Xu, C. & Jiang, X. MicroRNA-148a represents an independent prognostic marker in bladder cancer. Tumour biology: the journal of the International Society for Oncodevelopmental Biology and Medicine 37, 7915–7920, doi:10.1007/s13277-015-4688-0 (2016).

16 Bertorelle, R. et al. Telomerase is an independent prognostic marker of overall survival in patients with colorectal cancer. British journal of cancer 108, 278–284, doi:10.1038/bjc.2012.602 (2013).

17 Deng, F. et al. Overexpression of KIAA1199: An independent prognostic marker in nonsmall cell lung cancer. Journal of cancer research and therapeutics 13, 664–668, doi:10.4103/jcrt.JCRT_61_17 (2017).

18 Gu, L. et al. Cthrc1 overexpression is an independent prognostic marker in gastric cancer. Human pathology 45, 1031–1038, doi:10.1016/j.humpath.2013.12.020 (2014).

19 Vogelstein, B. et al. Cancer genome landscapes. Science 339, 1546–1558, doi:10.1126/science.1235122 (2013).

20 Samur, M. K. RTCGAToolbox: a new tool for exporting TCGA Firehose data. PloS one 9, e106397, doi:10.1371/journal.pone.0106397 (2014).

21 Stratford, J. K. et al. A six-gene signature predicts survival of patients with localized pancreatic ductal adenocarcinoma. PLoS medicine 7, e1000307, doi:10.1371/journal.pmed.1000307 (2010).

22 Leighton, J. C., Jr. & Goldstein, L. J. P-glycoprotein in adult solid tumors. Expression and prognostic significance. Hematology/oncology clinics of North America 9, 251–273 (1995).

23 Wang, Y., Goodison, S., Li, X. & Hu, H. Prognostic cancer gene signatures share common regulatory motifs. Scientific reports 7, 4750, doi:10.1038/s41598-017-05035-3 (2017).

24 Beyenbach, K. W. & Wieczorek, H. The V-type H+ ATPase: molecular structure and function, physiological roles and regulation. The Journal of experimental biology 209, 577–589, doi:10.1242/jeb.02014 (2006).

25 Zhang, X., Lv, Q. L., Huang, Y. T., Zhang, L. H. & Zhou, H. H. Akt/FoxM1 signaling pathway-mediated upregulation of MYBL2 promotes progression of human glioma. Journal of experimental & clinical cancer research: CR 36, 105, doi:10.1186/s13046-017-0573-6 (2017).

26 Guan, Z., Cheng, W., Huang, D. & Wei, A. High MYBL2 expression and transcription regulatory activity is associated with poor overall survival in patients with hepatocellular carcinoma. Current research in translational medicine 66, 27–32, doi:10.1016/j.retram.2017.11.002 (2018).

27 Fan, X. et al. B-Myb Mediates Proliferation and Migration of Non-Small-Cell Lung Cancer via Suppressing IGFBP3. International journal of molecular sciences 19, doi:10.3390/ijms19051479 (2018).

28 Hong, S. A. et al. Prognostic value of Dickkopf-1 and ss-catenin expression in advanced gastric cancer. BMC cancer 18, 506, doi:10.1186/s12885-018-4420-8 (2018).

29 Skiriute, D. et al. Promoter methylation of AREG, HOXA11, hMLH1, NDRG2, NPTX2 and Tes genes in glioblastoma. Journal of neuro-oncology 113, 441–449, doi:10.1007/s11060-013-1133-3 (2013).

30 Carlson, M. R. et al. Relationship between survival and edema in malignant gliomas: role of vascular endothelial growth factor and neuronal pentraxin 2. Clinical cancer research: an official journal of the American Association for Cancer Research 13, 2592–2598, doi:10.1158/1078-0432.CCR-06-2772 (2007).

31 Izawa, D. & Pines, J. How APC/C-Cdc20 changes its substrate specificity in mitosis. Nature cell biology 13, 223–233, doi:10.1038/ncb2165 (2011).

32 Gassmann, R. et al. Borealin: a novel chromosomal passenger required for stability of the bipolar mitotic spindle. The Journal of cell biology 166, 179–191, doi:10.1083/jcb.200404001 (2004).

33 Musa, J., Aynaud, M. M., Mirabeau, O., Delattre, O. & Grunewald, T. G. MYBL2 (B-Myb): a central regulator of cell proliferation, cell survival and differentiation involved in tumorigenesis. Cell death & disease 8, e2895, doi:10.1038/cddis.2017.244 (2017).

34 Diril, M. K. et al. Cyclin-dependent kinase 1 (Cdk1) is essential for cell division and suppression of DNA re-replication but not for liver regeneration. Proceedings of the National Academy of Sciences of the United States of America 109, 3826–3831, doi:10.1073/pnas.1115201109 (2012).

35 Zhu, C. et al. Functional analysis of human microtubule-based motor proteins, the kinesins and dyneins, in mitosis/cytokinesis using RNA interference. Molecular biology of the cell 16, 3187–3199, doi:10.1091/mbc.e05-02-0167 (2005).

36 Corson, T. W. et al. KIF14 messenger RNA expression is independently prognostic for outcome in lung cancer. Clinical cancer research: an official journal of the American Association for Cancer Research 13, 3229–3234, doi:10.1158/1078-0432.CCR-07-0393 (2007).

37 Stanietsky, N. et al. The interaction of TIGIT with PVR and PVRL2 inhibits human NK cell cytotoxicity. Proceedings of the National Academy of Sciences of the United States of America 106, 17858–17863, doi:10.1073/pnas.0903474106 (2009).

38 Baumann, C., Korner, R., Hofmann, K. & Nigg, E. A. PICH, a centromere-associated SNF2 family ATPase, is regulated by Plk1 and required for the spindle checkpoint. Cell 128, 101–114, doi:10.1016/j.cell.2006.11.041 (2007).

39 Jiang, K. et al. Microtubule minus-end regulation at spindle poles by an ASPM-katanin complex. Nature cell biology 19, 480–492, doi:10.1038/ncb3511 (2017).

40 Spiegel, I., Salomon, D., Erne, B., Schaeren-Wiemers, N. & Peles, E. Caspr3 and caspr4, two novel members of the caspr family are expressed in the nervous system and interact with PDZ domains. Molecular and cellular neurosciences 20, 283–297 (2002).

41 Xu, T. et al. High G2 and S-phase expressed 1 expression promotes acral melanoma progression and correlates with poor clinical prognosis. Cancer science 109, 1787–1798, doi:10.1111/cas.13607 (2018).

42 Sturm, D. et al. Hotspot mutations in H3F3A and IDH1 define distinct epigenetic and biological subgroups of glioblastoma. Cancer cell 22, 425–437, doi:10.1016/j.ccr.2012.08.024 (2012).

43 Dai, X. et al. Breast cancer intrinsic subtype classification, clinical use and future trends. American journal of cancer research 5, 2929–2943 (2015).

44 Cancer Genome Atlas Research, N. et al. Integrated genomic characterization of oesophageal carcinoma. Nature 541, 169–175, doi:10.1038/nature20805 (2017).

45 Morris, L. G. et al. Pan-cancer analysis of intratumor heterogeneity as a prognostic determinant of survival. Oncotarget 7, 10051–10063, doi:10.18632/oncotarget.7067 (2016).

46 An, N., Yu, Z. & Yang, X. Expression Differentiation Is Not Helpful in Identifying Prognostic Genes Based on TCGA Datasets. Molecular therapy. Nucleic acids 11, 292–299, doi:10.1016/j.omtn.2018.02.013 (2018).

47 Onuchic, L. F. et al. PKHD1, the polycystic kidney and hepatic disease 1 gene, encodes a novel large protein containing multiple immunoglobulin-like plexin-transcription-factor domains and parallel beta-helix 1 repeats. American journal of human genetics 70, 1305–1317, doi:10.1086/340448 (2002).

## References

1 Samur, M. K. RTCGAToolbox: a new tool for exporting TCGA Firehose data. PloS one 9, e106397, doi:10.1371/journal.pone.0106397 (2014).

2 Stratford, J. K. et al. A six-gene signature predicts survival of patients with localized pancreatic ductal adenocarcinoma. PLoS medicine 7, e1000307, doi:10.1371/journal.pmed.1000307 (2010).

3 Xu-Monette, Z. Y., Zhou, J. & Young, K. H. PD-1 expression and clinical PD-1 blockade in B-cell lymphomas. Blood 131, 68–83, doi:10.1182/blood-2017-07-740993 (2018).

4 Stanietsky, N. et al. The interaction of TIGIT with PVR and PVRL2 inhibits human NK cell cytotoxicity. Proceedings of the National Academy of Sciences of the United States of America 106, 17858–17863, doi:10.1073/pnas.0903474106 (2009).

5 Bailey, M. H. et al. Comprehensive Characterization of Cancer Driver Genes and Mutations. Cell 174, 1034–1035, doi:10.1016/j.cell.2018.07.034 (2018).

6 Lawrence, M. S. et al. Mutational heterogeneity in cancer and the search for new cancer-associated genes. Nature 499, 214–218, doi:10.1038/nature12213 (2013).

7 Huang da, W., Sherman, B. T. & Lempicki, R. A. Systematic and integrative analysis of large gene lists using DAVID bioinformatics resources. Nature protocols 4, 44–57, doi: 10.1038/nprot.2008.211 (2009).

8 Tarca, A. L. et al. A novel signaling pathway impact analysis. Bioinformatics 25, 75–82, doi: 10.1093/bioinformatics/btn577 (2009).

